# Sleep spindles preferentially consolidate weakly encoded memories

**DOI:** 10.1101/2020.04.03.022434

**Authors:** Dan Denis, Dimitrios Mylonas, Craig Poskanzer, Verda Bursal, Jessica D. Payne, Robert Stickgold

## Abstract

Sleep has been shown to be critical for memory consolidation, with some research suggesting that certain memories are prioritized for consolidation. Initial strength of a memory appears to be an important boundary condition in determining which memories are consolidated during sleep. However, the role of consolidation-mediating oscillations, such as sleep spindles and slow oscillations, in this preferential consolidation has not been explored. Here, 54 human participants (76% female) studied pairs of words to three distinct encoding strengths, with recall being tested immediately following learning and again six hours later. Thirty-six had a two-hour nap opportunity following learning, whilst the remaining 18 remained awake throughout. Results showed that across six hours awake, weakly encoded memories deteriorated the fastest. In the nap group however, this effect was attenuated, with forgetting rates equivalent across encoding strengths. Within the nap group, consolidation of weakly encoded items was associated with fast sleep spindle density during non-rapid eye movement sleep. Moreover, sleep spindles that were coupled to slow oscillations predicted the consolidation of weak memories independently of uncoupled sleep spindles. These relationships were unique to weakly encoded items, with spindles not correlating with memory for intermediate or strong items. This suggests that sleep spindles facilitate memory consolidation, guided in part by memory strength.

**Significance statement:** Given the countless pieces of information we encode each day, how does the brain select which memories to commit to long-term storage? Sleep is known to aid in memory consolidation, and it appears that certain memories are prioritized to receive this benefit. Here, we found that compared to staying awake, sleep was associated with better memory for weakly encoded information. This suggests that sleep helps attenuate the forgetting of weak memory traces. Fast sleep spindles, a hallmark oscillation of non-rapid eye movement sleep, mediate consolidation processes. We extend this to show that fast spindles were uniquely associated with the consolidation of weakly encoded memories. This provides new evidence for preferential sleep-based consolidation and elucidate a physiological correlate of this benefit.

## Introduction

Sleep aids in the consolidation of memories (Stickgold, 2005; Payne et al., 2008a; Klinzing et al., 2019). Some evidence suggests that this process is selective, with certain memories being prioritized for retention over others (Payne et al., 2008b; Diekelmann et al., 2009; Stickgold and Walker, 2013; Payne and Kensinger, 2018). It appears that certain salience cues, such as emotional valence or personal relevance, present during the peri-encoding period can act as behavioral ‘tags’ that indicate which memories should be consolidated during sleep (Payne et al., 2008b, 2012, 2015; Fischer and Born, 2009; Wilhelm et al., 2011). Tagging of memories during encoding may be realized through changes in arousal-related neuromodulators such as norepinephrine and cortisol, and functional brain connectivity (Kim and Payne, 2020). The initial strength of a memory appears to act as a boundary condition of sleep-based consolidation, but the direction of the effect is unclear (Tucker and Fishbein, 2008; Schapiro et al., 2017; Schoch et al., 2017; Denis et al., 2020)

Several studies have manipulated encoding strength by varying the number of item presentations given during encoding. Using this method, many of these studies have found that *weaker* memories are prioritized for consolidation (Drosopoulos et al., 2007; Schapiro et al., 2017; Denis et al., 2020). Increasing neural pattern similarity has been seen across successive item presentations and is associated with better memory (Xue et al., 2010; Lu et al., 2015), and a large number of repetitions leads to stronger cortical memory representations (Brodt et al., 2018). It is thus possible that, after multiple presentations, memory traces have already been rendered strong enough that sleep does not or cannot strengthen them further, resulting in the sleep benefit being strongest for initially weak memories. It appears that a minimum threshold does need to be met however, such as a certain degree of hippocampal recruitment during encoding (Rauchs et al., 2011), the utilization of deep encoding strategies such as successful visualization (Denis et al., 2020), or successful recall during a pre-sleep memory test (Denis et al., 2020; Muehlroth et al., 2020). When weak memories are formed due to acute sleep restriction, sleep has been shown to rescue these memories, strengthening the neural representation across a night of recovery sleep (Baena et al., 2020).

On the other hand, others have reported that strong memories are prioritized. Some work has concluded this on the basis of only the top half of initial learners benefitting from sleep (Tucker and Fishbein, 2008; Wislowska et al., 2017). However this may conflate weak vs strong encoding with good vs poor learners, which may not tap into the same process (Creery et al., 2015). The act of memory retrieval can also strengthen memories (Roediger and Butler, 2011). It has been suggested that a post-encoding pre-sleep retrieval test is necessary for sleep-based consolidation to be detected (Schoch et al., 2017). Other work utilizing a series of behavioral studies reported no benefit of sleep following post-encoding, pre-sleep memory retrieval without feedback (Bäuml et al., 2014; Abel et al., 2019). This finding was interpreted in relation to a bifurcation model, whereby successful retrieval strengthens items to such an extent that sleep does not need to consolidate them, whereas failed retrievals are too weak to benefit (Bäuml et al., 2014). This is conceptually similar to the proposal that memories are consolidated during sleep based on an inverted u-shape distribution, with memories closer to the middle of the strength scale receiving the largest benefit (Stickgold, 2009).

There is little work on the sleep correlates of this consolidation benefit based on initial encoding strength. One study reported that the benefit of a nap for weakly encoded items was associated with both non-rapid eye movement (NREM) and rapid eye movement (REM) sleep, with NREM followed by a larger amounts of REM sleep being optimal for preferential memory consolidation (Schapiro et al., 2017). The active systems consolidation theory of memory consolidation posits that during NREM sleep, memories supported by the hippocampus are repeatedly reactivated through the triple phase-locking of hippocampal sharp-wave ripples, thalamocortical sleep spindles, and neocortical slow oscillations (Rasch and Born, 2013; Staresina et al., 2015). Specifically, de-polarizing slow oscillation upstates are thought to facilitate the emergence of sleep spindles, which in turn mediate transfer of information reactivated during sharp wave ripples in the hippocampus, leading to long-term storage more dependent on neocortical sites (Klinzing et al., 2019). There is evidence for these oscillations, and especially their coupling, being involved in general memory consolidation processes (Niknazar et al., 2015; Latchoumane et al., 2017; Mikutta et al., 2019; Muehlroth et al., 2019; Cross et al., 2020).

It appears that two distinct types of spindle exist, fast and slow frequency types (Cox et al., 2017; Fernandez and Luthi, 2020), and it has been observed that their coupling to the slow oscillations differ (Mölle et al., 2011; Cox et al., 2014, 2018; Klinzing et al., 2016). In young healthy adults, fast spindles appear to couple just before or at the positive peak of the slow oscillation (Cox et al., 2018; Helfrich et al., 2018). Slow spindles, on the other hand, preferentially couple in the transition from the positive peak to the trough (Cox et al., 2018). These differences in coupling behavior may translate to different functional roles for fast and slow spindle types (Schabus et al., 2006; Lustenberger et al., 2015; Cowan et al., 2020; Fernandez and Luthi, 2020).

Recent work has demonstrated coupling can rescue poorly formed memories formed following sleep restriction (Baena et al., 2020), but it is currently unknown if these oscillations act preferentially based on the encoding strength of a memory when memory strength is manipulated within-participants under more typical encoding conditions, and whether fast or slow spindles are preferentially involved.

In a typical overnight design, the wake control group will learn information in the morning and be tested in the evening, whereas the sleep group learns in the evening and is tested the following morning. A nap design, in addition to being simpler to implement, allows learning and test phases to occur at the same time of day for all participants, eliminating possible circadian confounds and making the role of sleep clearer (Mednick et al., 2003; Payne et al., 2005, 2009; Lo et al., 2014; c.f. Schalkwijk et al., 2019). It also restricts the amount of time spent awake that might expose participants to interfering information.

In this study, we used a daytime nap to investigate unresolved questions regarding the role of encoding strength in sleep-based memory consolidation. Participants spent the day in the sleep laboratory. In the morning, they learned word pairs to differing levels of encoding strength. Some participants then had a two-hour nap opportunity and were tested on their memory 4 hours later. Other participants remained awake in the lab throughout. We sought to understand 1) whether a nap prioritizes the consolidation of memories based on their encoding strength in a similar manner to a full night of sleep; and 2) whether sleep oscillations (namely sleep spindles and their coupling with slow oscillations) facilitate preferential consolidation.

## Methods

### Participants

In total, 54 human participants completed the full study protocol. The mean age of participants was 22 (SD = 3) years, and 76% were female. Participants reported no history of any sleep, neurological, or psychiatric disorders, normal bedtimes no later than 2am, and sleeping on average for at least six hours each night. For the three days prior to the study, participants were instructed to keep to a regular sleep schedule and abstain from caffeine on the morning of the study. Recruitment was through advertisements for a study of learning and memory placed on local college job boards. Participants received financial compensation for their time. The study received IRB approval from Beth Israel Deaconess Medical Center.

### Design

The study design is depicted in **Figure 1**. All participants followed the same experimental procedure, except for whether they were allowed to take a nap (**Figure 1A**). After providing informed consent, participants filled out questionnaires about their subjective sleep habits over the past three days, their general sleep quality over the past month (assessed with the Pittsburgh Sleep Quality Index (PSQI; Buysse et al., 1989), and their current subjective sleepiness and alertness levels (Stanford Sleepiness Scale (SSS; Hoddes et al., 1972). No actigraphy was collected. Following this, participants were wired for EEG (see below). Then, they took part in the first experimental session. The session started with a 5-minute eyes-closed quiet rest session (all subsequent rest sessions were also 5 minutes eyes closed and will be analyzed in future studies of resting state activity). They then studied pairs of words and were asked to try and visualize a scene containing the two objects described by the word pair (**Figure 1B**). After encoding, participants had a second quiet rest session, and then a cued recall test (immediate recall; **Figure 1C**), and finally a third rest period. After this, 36 participants were told they now have a 2-hour opportunity to nap, which was then followed by four hours spent awake in the lab watching TV. The remaining 18 participants were not given the opportunity to nap, so remained awake in the lab for six hours. These group sizes were similar to our previous publication that successfully demonstrated preferential consolidation of weakly encoded material (Denis et al., 2020). The nap group was oversampled due to our interest in sleep-physiological activity *within* this group. The second experimental session occurred after the six-hour delay period. During this session, participants had a fourth quiet rest period, followed by a second cued recall test (delayed recall; **Figure 1C**), and a fifth and final quiet rest session. Finally, at the very end of the protocol, participants filled out two additional questionnaires assessing trait abilities in forming internal visualizations [measured using the vividness of visual imagery questionnaire (VVIQ; Marks, 1973) and the visual portion of the Plymouth sensory imagery questionnaire (PSIQ; Andrade et al., 2014)].

**Figure 1.**
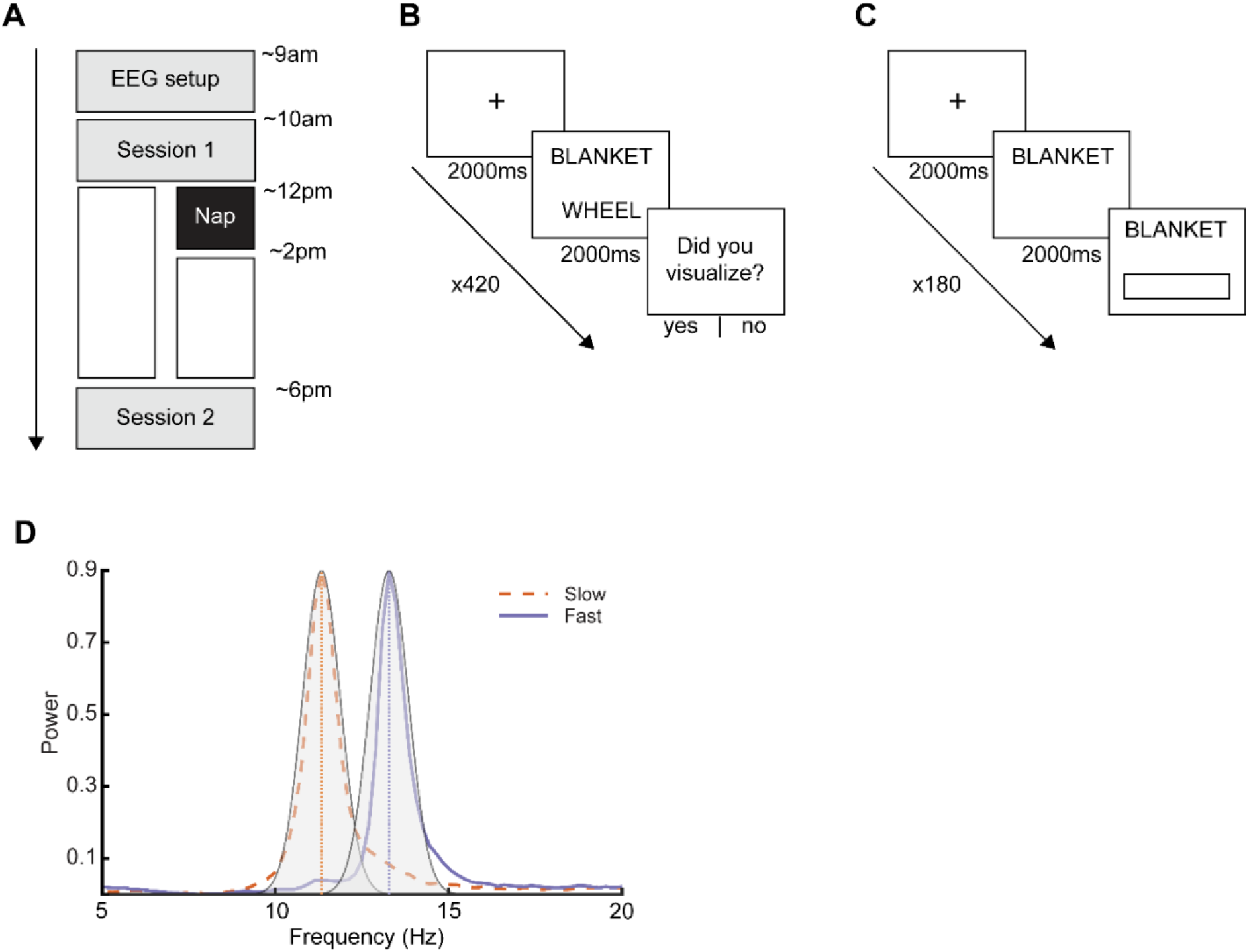
Experimental design. **A** – Timeline of the protocol. All participants arrived at the sleep laboratory at ~9am and were wired for EEG. During Session 1, participants completed a 5-minute rest period followed by the encoding task (**B**), and a second 5-minute rest session. They then performed a cued recall test (immediate recall; **C**), followed by a final quiet rest session. Following that, 36 participants had a two-hour nap opportunity, followed by a four-hour wake delay spent in the sleep laboratory. The remaining 18 did not have a nap opportunity and stayed awake in the lab for six hours. Session 2 started ~6pm and began with a fourth quiet rest, followed by a second cued recall test (delayed recall; **C**), and a fifth quiet rest. **B** – Encoding. Each encoding trial began with a fixation cross that appeared on the screen for 2000-3000ms, followed by the word pair for 2000ms. After the presentation of each word pair, participants were asked if they had been able to successfully visualize a scene containing the two word-pair objects. A total of 180-word pairs were displayed, with 60 being viewed 1 time, 60 being viewed 2 times, and 60 being viewed 4 times, for a total of 420 trials. **C** – Recall. Both the immediate and delayed test followed the same procedure. First, a fixation cross appeared for 2000-3000ms. Then, the first word of the pair appeared alone for 2000-2500ms. During this window, participants were instructed to think as hard as possible about what the correct second word was. Then, a box appeared underneath the first word, indicating that they could start typing in their answer. There was a total of 180 recall trials. **D** – Purple and orange lines show fast and slow spindle components as derived using generalized eigen decomposition from a single participant. Overlaid in gray is the frequency response of the wavelet used for spindle detection, separately tuned for fast and slow spindles based on the participant’s peaks. Note the low degree of overlap in the fast and slow wavelet frequency response.

### Encoding

During the encoding task, participants studied 180 pairs of words. Participants were instructed, for each trial, to try to visualize a scene containing the two objects in the word pair (*e.g*., “blanket – wheel”). Word pairs were assigned to either a weak (n = 60), intermediate (n = 60), or strong (n = 60) encoding condition, with assignments randomized across participants. Word pairs in the weak condition were presented once (n = 60 trials), pairs in the intermediate condition twice (n = 120 trials), and those in the strong condition four times (n = 240 trials), for a total of 420 trials. In a prior study, we demonstrated that this procedure produces distinct levels of encoding strength (Denis et al., 2020). The order of presentation was pseudorandomized for each participant, with at least two trials separating multiple presentations of any one item.

On each trial (**Figure 1B**), a fixation cross appeared in the center of the screen for 2,000-3,000ms, followed by the word pair for 2,000ms. This was followed by a blank screen for 500ms-1,000ms. Participants were then asked whether they had visualized a scene containing both objects, responding (either yes or no) by a keypress. After responding, a blank screen appeared for 1,000ms, and then the next trial began. After every 70 trials, there was a break lasting a minimum of one minute and terminated by the participant. The variation in presentation times for the fixation cross and blank screen was to facilitate future event-related EEG analyses on the encoding data, with the jitter allowing for the best assessment of memory encoding activity, rather than preparatory responses to the stimuli.

### Recall

Both the immediate and delayed recall tests followed the same procedure (**Figure 1C**). Each trial began with a fixation cross on the screen for 2,000ms-3,000ms. Then, the first word of the pair appeared for 2,000ms. During this period, participants were instructed to recall the second word of the pair, but not to type it. After 2000-2500ms, a box appeared under the first word, indicating participants could enter their answer. This approach allowed for time-locked analysis of memory recall in future analyses. A separate study (n = 52) confirmed that variable presentation times in this window did not impact immediate memory performance (not reported). Participants were instructed to respond as quickly and as accurately as possible, and that there was no penalty for guessing. If no response was entered after seven seconds, a prompt appeared telling participants to respond, and if the participant had not begun typing a response after a further three seconds, the program advanced to the next trial. Each word pair was tested once, for a total of 180 trials. Order of presentation of the word pairs was randomized for each session and for each participant. At the end of the immediate recall session, participants were told that their memory for the word pairs would be tested again at the end of the day. All tasks were administered using custom scripts written in the Psychtoolbox package for MATLAB (Kleiner et al., 2007).

### EEG acquisition and preprocessing

EEG was collected from all participants throughout the protocol. During the delay period, participants remained connected to the EEG equipment, but no data were acquired. Only EEG data recorded during the nap is reported here. Data were acquired from 57 EEG channels, with electrodes positioned in accordance with the 10-20 system. Additionally, electrodes were placed on the left and right mastoids, above the right eye and below the left eye (for EOG measurements), two placed on the chin (for EMG measurements), one on the forehead (recording reference) and one on the collarbone (ground). An Aura-LTM64 amplifier and TWin software were used for data acquisition (Grass Technologies, Warwick, RI). All impedances were kept below 25 KOhm. The sampling rate was 400Hz.

Sleep scoring was performed according to standard criteria (Iber et al., 2007). Sleep scoring and subsequent sleep statistic generation was performed in MATLAB (The Mathworks, Natick, MA). EEG analyses were performed on the full high-density EEG array using custom MATLAB scripts. First, all EEG channels were re-referenced to the average of the two mastoids, band-pass filtered between 0.3 - 35Hz and notch filtered at 60Hz. Data was then artifact rejected based on visual inspection of each 30-second epoch. Bad epochs were marked and removed from subsequent analyses, and bad channels were marked and interpolated using a spherical splines algorithm implemented in EEGLAB (Delorme and Makeig, 2004). All artifact-free data were then subjected to further analysis.

### Sleep spindle detection

Spindles were automatically detected at every electrode during non-rapid eye movement sleep (NREM; Stage 2 + SWS) sleep using a modification of a previously validated detector to account for inter-individual differences in the peak frequency of slow and fast spindles (Wamsley et al., 2012; Mylonas et al., 2019). As a first step, slow and fast spindle peaks were identified through inspection of the NREM sleep power spectrum. Because it can be difficult to detect a slow spindle peak in many individuals by looking at the channel-level spectrum, we instead utilized generalized eigendecomposition (GED) to maximally separate slow and fast spindle activity (Cohen, 2017; Cox et al., 2017). In brief, GED operates on two separate covariance matrices to find eigenvectors that maximally differentiate the case. We constructed one covariance matrix from the NREM time series filtered in a broad slow spindle range (9-12.5Hz), and the second covariance matrix from the NREM time series filtered in the fast spindle range (12.5-16Hz). High order (13,200) filters were applied to ensure minimal overlap between slow and fast spindle ranges. This yielded a fast and a slow covariance matrix (57 × 57 channels) from which we generated a 57 × 57 matrix of eigenvectors. The eigenvector with the highest eigenvalue maximizes slow relative to fast spindle power, and conversely, the eigenvector with the lowest eigenvalue maximizes fast relative to slow spindle power (Cox et al, 2017).

Next, we multiplied the full, raw EEG time series with the full eigenvector matrix, resulting in a time series of 57 components (Cox et al., 2017). This component time series was then transformed to the frequency domain by estimating the power spectral density (PSD) of the derivative of the time series using Welch’s method with 5 second windows and 50% overlap. PSD was estimated on the derivative of the time series, as opposed to the time series itself, to minimize 1/*f* scaling and make spectral peaks in the data easier to identify (Sleigh et al., 2001; Demanuele et al., 2007). The resulting component spectra were then visualized (the first four (for slow spindles) and last four (for fast spindles) were plotted), and each participant’s slow and fast spindle peak were manually identified. This approach yielded an average slow spindle peak frequency of 11.04Hz (SD 0.92Hz, range = 9.18Hz – 12.50Hz) and an average fast spindle peak frequency of 13.81Hz (SD = 0.57Hz, range = 12.70Hz – 14.84Hz).

After detecting individuals’ slow and fast spindle peak frequencies, sleep spindles were automatically detected using a wavelet-based detector. The raw EEG signal was subjected to a time-frequency transformation using complex Morlet wavelets. The wavelet parameters were tuned for each individual based on their slow and fast spindle peaks. Specifically, the peak frequency of the wavelet was set at that individual’s slow or fast spindle peak, with the bandwidth of the wavelet (full-width half-max) being a 1.3Hz range centered on the peak frequency (e.g. if a participant’s fast spindle peak was 13Hz, the wavelet peak frequency would be set at 13Hz, with a bandwidth of 12.35-13.65Hz). This narrow bandwidth was motivated in order to minimize overlap between the fast and slow spindle ranges (**Figure 1D**), and is consistent with prior research (Cox et al., 2017). Spindles were detected on each channel by applying a thresholding algorithm to the extracted wavelet scale. A spindle was detected whenever the wavelet signal exceeded a threshold of 6 times the median signal amplitude of all artifact-free epochs for a minimum of 400ms. The threshold of 6 times the median was determined empirically for both slow and fast spindles, using Otsu’s method to maximize between-class (‘spindle’,’non-spindle’) variance in the wavelet coefficient (Otsu, 1979, Djonlagic et al., 2020). Our main metric of focus was spindle density (spindles per minute) in NREM sleep.

### Slow oscillation detection

Slow oscillations were detected at every electrode during NREM sleep using a second automated algorithm that band-pass filtered the EEG between 0.5 and 4Hz and identified all positive-to-negative zero crossings (Staresina et al., 2015; Helfrich et al., 2018). Candidate slow oscillations were marked if two such consecutive zero crossings fell 0.5 – 2.0 seconds apart. Peak-to-peak amplitudes for all candidate slow oscillations were determined, and oscillations in the top quartile (*i.e*., with the highest amplitudes) at each electrode were retained as slow oscillations. The use of this cutoff has been used in previous research (Staresina et al., 2015; Helfrich et al., 2018).

### Slow oscillation-spindle coupling

Slow oscillation-spindle coupling were identified at every electrode during NREM sleep, separately for slow and fast spindles. First, EEG data was band-pass filtered in the delta (0.5 −4Hz), and each participant’s slow and fast spindle band (again using a 1.3Hz bandpass centered around peak slow or fast spindle frequency). Then, the Hilbert transform was applied to extract the instantaneous phase of the delta-filtered signal and instantaneous amplitude of the spindle-filtered signal. For each detected spindle, the peak amplitude of that spindle was determined. It was then determined whether the spindle peak occurred within the time course (i.e. between the two positive-to-negative crossings) of any detected slow oscillation. If the spindle peak was found to occur during a slow oscillation, the phase angle of the slow oscillation at the peak of the spindle was determined. Finally, for each electrode on each participant, we extracted the percentage of all spindles that were coupled with slow oscillations, the coupled and uncoupled spindle densities, and the average coupling phase angle for all coupled spindles. Coupling strength was assessed as the mean vector length.

Additional visualization methods were employed to better display the temporal dynamics of slow oscillation-spindle coupling events in order to increase our confidence that fast and slow spindle types had been reliably separated. First, peri-event histograms were calculated to illustrate the timing of fast and slow spindle couplings to the slow oscillation. These histograms express the distribution of coupling events (% of spindles coupled, averaged across participants) in a −1000 – 1000ms window centered on the trough of the slow oscillation (t0), in 100ms bins. Separate histograms were constructed for fast and slow spindle coupling. Second, time-frequency representations of coupling dynamics were calculated. For this, we first located all the slow oscillations coupled to either a fast or a slow spindle. Data were then epoched, separately for fast and slow coupling events, into −3000 – 3000ms trials centered on the trough of the coupled slow oscillations (t0). Time frequency analysis was applied using complex Morlet wavelets ranging from 2 – 30Hz (40 frequencies, wavelet cycles increasing linearly from 4 – 12 cycles). Power was decibel normalized (10 * log10(power / baseline), where the baseline was average power in a =1500-1500ms period centered on the slow oscillation trough (Kurz et al., 2020).

To better assess whether coupling results reflect a “true”co-occurrence of the two oscillations, we needed to ensure that the number of coupling events exceeded what would be expected by chance, given the number of detected slow oscillations, sleep spindles, and time spent in NREM sleep. To this end, the observed signal was compared to a randomized signal where the spindle signal was circularly shifted, and coupling was recalculated. Circular shifting is a widely used method to create surrogate data for nonparametric tests (Gilson et al., 2017). The new start of the signal is set to a random position and early entries are sequentially moved to after the end of the original signal. By shifting the time series in this fashion (in this case the spindle signal) the temporal relationship between spindles and slow oscillations is disrupted, whereas the signal properties under consideration (i.e. the distribution of spindles and slow oscillations) are retained. This was performed over 1,000 iterations to generate a null distribution of slow oscillation-spindle coupling. The null distribution was then compared to the observed values. Across each participant/electrode, we calculated the percentage of participants and electrodes where the degree of coupling was significantly higher (*p* < .05) than what would be expected by chance.

### Statistical analysis

Behavioral data were assessed using linear mixed effect models, implemented in the *lme4* package for R (Bates et al., 2015; R Core Team, 2018). As fixed effects, we entered the interaction between item presentation condition and group. Participant was entered as a random intercept:

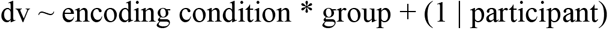

where the dv was either: Percentage of items visualized, percentage of items recalled at immediate test, percentage of items recalled at delayed test, or the relative change in percentage recall between tests [(% delayed recall - % immediate recall) / %immediate recall].

Statistical significance was obtained by likelihood ratio tests (LRT) with the full model (encoding condition * group interaction) compared to reduced models containing main effects only. A statistically significant LRT indicates that the full model is a better fit than the sub-model in question, whilst a non-significant LRT suggests the sub-model to be more appropriate. In cases where multiple models were significant/non-significant, the most parsimonious model was selected. After obtaining the most optimal model, pairwise contrasts were performed where appropriate with the emmeans package for R (Lenth, 2018). These contrasts produced the estimate and 95% confidence intervals for each pairwise contrast.The false discovery rate (FDR) was used to adjust *p* values for multiple pairwise comparisons.

We correlated change in memory against sleep oscillatory EEG measures at each electrode. To control for multiple comparisons across electrodes, and to take into account the spatial correlation of the high density EEG data, we used a cluster-based permutation approach in the FieldTrip toolbox for MATLAB, using the *ft_statfun_correlationT* function (Oostenveld et al., 2011). All analyses used the following parameters: 10,000 iterations; a *clusteralpha* of 0.05 with the default *maxsum* method to determine cluster significance; and a significant threshold of 0.05. For comparisons between fast and slow spindle properties, the same procedure was followed except the *ft_statfun_depsamplesT* function was used.

To better visualize significant correlations, scatterplots were also created. For these, activity at each electrode that was significant in the overall cluster was averaged together for the purposes of visualization in the scatterplot. In order to minimize the influence of outliers, these associations were assessed using robust regression procedures (MATLAB function *lmfit* with *RobustOptions* turned on). Correlation coefficients was used to obtain a standard estimate of association strength, which are not provided by the robust regression procedures. Robust regressions were performed on these averaged data rather than scalp-wide cluster-based test because robust regression procedures are not implemented in the FieldTrip environment.

Comparisons between fast and slow spindle coupling phase were performed using the Hotelling paired sample test for circular means (van den Brink et al., 2014; van den Brink, 2020). Correlations between coupling phase angle and memory were performed using circular-linear correlations, implemented in the CircStat toolbox for MATLAB (Berens, 2009). As circular analyses are not implemented in the FieldTrip environment, the FDR was used to control for multiple comparisons. Comparisons of correlation coefficient magnitudes were performed using Meng’s Z test for within-group comparisons (Meng et al., 1992; Spaak, 2020). Robust multiple linear regression was used to assess the independent contribution of coupled and uncoupled spindles to changes in memory across the delay interval. The independent contributions of each predictor, while controlling for the other predictor, were then visualized using partial regression plots to aid interpretation of the multiple regression (Velleman and Welsch, 1981). These scatterplots depict the relationship between the dependent variable (*y*) and one of the predictors (*x_0_*) in a multiple regression model, after controlling for the presence of the other predictor (*x_1_*) (Velleman and Welsch, 1981). To create the plot, first *y* is regressed on *x_1_*, omitting *x_o_*. Second, *x_o_* is regressed on *x*_1_. By plotting the residuals of the first regression against the residuals of the second regression, it is possible to visually display the relationship between *x_0_* and *y* after controlling for variable *x_1_* (Velleman and Welsch, 1981).

## Results

### Behavior

Participants were highly successful at visualizing the word pairs, with 78% (SD = 21%) of the word pairs that were seen just once being reported as successfully visualized in a scene containing the two objects. We were therefore unable to look at differences in recall and consolidation between successfully and unsuccessfully visualized items, due to a lack of not-visualized trials. As such, all subsequent analyses report on all trials together.

We next looked at immediate recall accuracy to confirm that the presentation number manipulation successfully induced different levels of encoding strength (**Figure 2A**). Mean behavioral values can be found in **Table 1**. There was a significant main effect of presentation number on immediate recall performance (χ^2^ (4) = 149.50, *p* < .001), with significant increases in the percentage of word pairs recalled as the number of presentations during encoding increased (1PRES vs 2PRES = 27% increase, 1PRES vs 4RPES = 46% increase, 2PRES vs 4PRES = 19% increase; all *p* < .001). There was no difference between the groups (χ^2^ (3) = 0.81, *p* = .85), and there was no interaction between presentation number and group (χ^2^ (2) = 0.80, *p* = .67). These results show that the items were encoded at three distinct strengths, and that learning was equivalent between the nap and wake groups. When delayed recall accuracy was assessed (**Figure 2B**), the same pattern of results emerged (main effect of presentation number: χ^2^ (4) = 141.55, *p* < .001; main effect of group: χ^2^ (3) =1.43, *p* = .70; interaction: χ^2^ (2) = 1.27, *p* = .53).

**Figure 2.**
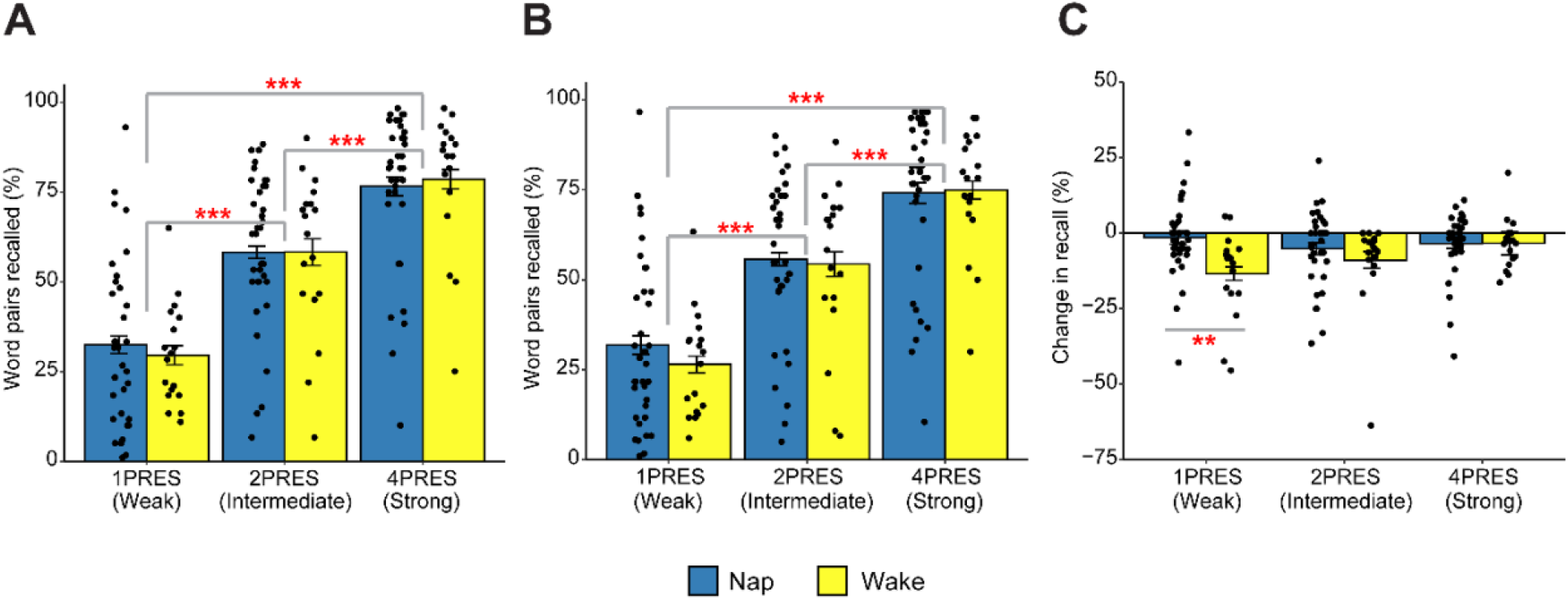
Behavior. **A** – Percentage of word pairs recalled during immediate test. **B** – Percentage of word pairs recalled during delayed test. **C** – Relative change in the percentage of word pairs recalled between delayed and immediate test. Note the change is calculated as %delayed – %immediate test / each individual participant’s %immediate test score. Error bars display the within-participant standard error. n PRES” indicates encoding condition e.g. 4PRES = recall/change in recall to items that were presented four times during encoding *** = *p* < .001, ** = *p* < .01

**Table 1.** Behavioral response data

|  | Immediate recall |  |  | Delayed recall |  |  | Delayed - immediate |  |  |
| --- | --- | --- | --- | --- | --- | --- | --- | --- | --- |
|  | 1PRES<br>M (SD) | 2PRES<br>M (SD) | 4PRES<br>M (SD) | 1PRES<br>M<br>(SD) | 2PRES<br>M<br>(SD) | 4PRES<br>M<br>(SD) | 1PRES<br>M<br>(SD) | 2PRES<br>M<br>(SD) | 4PRES<br>M<br>(SD) |
| Nap | 32.41<br>(22.82) | 58.24<br>(21.83) | 76.58<br>(21.29) | 31.87<br>(23.27) | 55.77<br>(22.15) | 74.16<br>(22.87) | -0.54<br>(3.05) | -2.47<br>(7.05) | -2.42<br>(7.52) |
| Wake | 29.57<br>(23.04) | 58.26<br>(21.77) | 78.61<br>(21.34) | 26.50<br>(23.50) | 54.35<br>(22.05) | 74.98<br>(22.90) | -3.07<br>(2.87) | -3.91<br>(7.04) | -3.63<br>(7.49) |

*Note.* 1PRES = Items presented once during encoding, 2 PRES = Items presented twice during encoding, 4PRES = Items presented four times during encoding

We then looked at the effects of a nap on memory at re-rest 6 hours later. We calculated, for each participant, the relative change in recall at delayed test compared to each participant’s immediate test score (**Figure 2C**). There was a significant main effect of group (χ^2^ (3) = 12.50, *p* = .006), with the nap group showing overall less forgetting (M = −3.20%, SD = 7.04%) than the wake group (M = −9.12%, SD = 14.57%). This suggests that across all items, sleep benefitted memory. There was no main effect of presentation number (χ^2^ (4) = 8.10,*p* = .088). There was, however, an interaction between presentation number and group (χ^2^ (2) = 6.37, *p* = .041), suggesting that the benefit of sleep on memory differed depending on encoding strength. Follow-up tests comparing relative change in recall between the nap and the wake group at each presentation number condition revealed a significant difference for 1PRES items (B [95% CI] = 11.96 [3.48, 20.400], *p* = .002). There was significantly more forgetting of 1PRES items in the wake group (M = −13.48%, SD = 13.84%) compared to the nap group (M = −1.53%, SD = 13.02%). The between group difference was not significant for either 2PRES (B [95% CI] = 3.71 [−4.77, 12.19], *p* = .44) or 4PRES (B [95%CI] = −0.06 [−8.54, 8.42], *p* = .99) items. Within the wake group, there was significantly more forgetting of 1PRES items than of 4PRES (M (SD) = −3.38% (8.23%)) items (B [95% CI] = −10.10 [−19.80, −0.41], *p* = .038). No other comparisons were significant (all *p* = .25). In the group which napped however, this increased forgetting of 1PRES items was eliminated, with no difference in the amount of forgetting between presentation number conditions (all *p* = .54).

Finally, we investigated whether the magnitude of the preferential memory consolidation effect differed between a nap and a full night of sleep, by comparing the present results to one of our previous datasets using a similar task where three encoding strengths were induced via the same item repetition procedure (Denis et al., 2020; **Figure 3**). Although the participants were different between studies, they were derived from the same population. Across wake only delays, forgetting of 1PRES items was significantly higher across a 12-hour waking delay (M = −31.62%, SD = 30.81%) compared to a 6-hour waking delay (M = −13.48%, SD = 13.84%), *t* (21.9) = 2.22, *p* = .037, *d* = 0.76. Across sleep however, forgetting was equivalent across a 12-hour delay containing nocturnal sleep (M = 0.19%, SD = 12.97%), and a 6-hour delay containing a 2-hour daytime nap opportunity (M = −1.53%, SD = 13.02%), *t* (34.2) = −0.46, *p* = .65, *d* = 0.13. This translated into a larger sleep-wake group difference across 12 hours (*d* = 1.35), than across 6 hours (*d* = 0.89). After a 24 hour delay (overnight sleep plus a day of wake), forgetting of 1PRES items was reduced compared to 12hours of wake only (M = −13.62%, SD = 14.36%), becoming comparable to the 6-hour wake delay (*t* (35) = 0.03, *p* = .98, *d* = 0.01).

**Figure 3.**
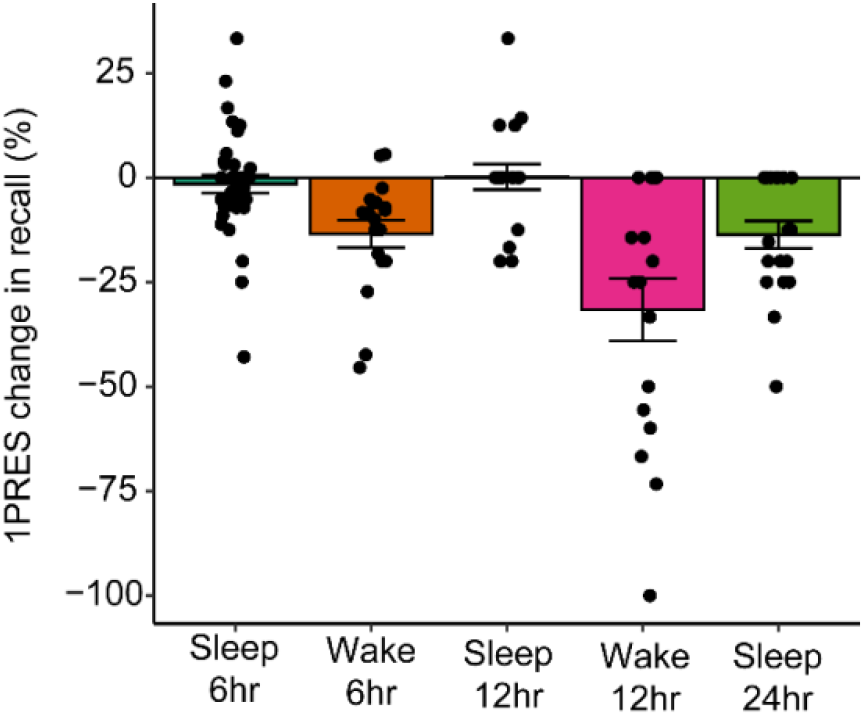
Change in 1PRES memory across delay periods. Note the data for the 12- and 24-hour delay groups have been previously published elsewhere (Denis et al, 2020).

The behavioral results show that a 2-hour nap opportunity significantly reduced forgetting over a 6-hour retention interval compared to staying awake. Over a wake only delay, items that were weakly encoded were forgotten at a higher rate than items that were more strongly encoded. After a nap + wake delay however, weakly encoded information showed similar retention to more strongly encoded information.

### Sleep architecture and sleep stage correlations

Sleep statistics are presented in **Table 2**. There were no significant correlations between change in memory (for either all items or any of the three encoding strengths) and time or percentage of the nap spent in any sleep stage (all *p*’s > .10).

**Table 2.** Sleep statistics

|  | Mean | SD |
| --- | --- | --- |
| Total sleep time (min) | 94.5 | 23.7 |
| Sleep onset latency (min) | 10 | 6.9 |
| Wake after sleep onset (min) | 12.2 | 18.1 |
| Sleep efficiency (%) | 79.5 | 17.1 |
| Stage 1 time (min) | 4.7 | 2.9 |
| Stage 2 time (min) | 48.5 | 17.1 |
| SWS time (min) | 21.6 | 14.4 |
| REM time (min) | 19.8 | 12.9 |
| Movement time (min) | 1.7 | 1.8 |
| Stage 1 percentage (%) | 5.1 | 4.0 |
| Stage 2 percentage (%) | 52.2 | 14.3 |
| SWS percentage (%) | 22.4 | 13.5 |
| REM percentage (%) | 20.2 | 12.1 |
| Movement percentage (%) | 1.9 | 2.1 |

*Note.* SD = Standard deviation. SWS = Slow wave sleep, REM = Rapid eye movement

### Sleep spindles

Topography of NREM detected fast and slow spindles are shown in **Figure 4A-B**. Fast spindle density exhibited a maxima at central electrode sites (M (SD) = 7.90 (1.62), range = 3.70 – 11.36, as measured at electrode Cz), whereas slow spindle density showed maximum activity in frontal regions (M (SD) = 5.19 (1.81), range = 2.13 – 9.14, as measured at electrode Fz). Fast spindle density was significantly higher at all electrodes compared to slow spindle density (cluster *t* = 443, *p* < .001). We calculated scalp-wide correlation coefficients between spindle density and change in recall for each of the encoding strengths. For fast spindle density, significant positive correlations were observed with change in memory for 1PRES items (cluster t = 110.64, *p* = .015; **Figure 4A**), with an average within-cluster correlation coefficient of *r* = .50 (**Figure 5**). This association was unique to the 1PRES items, with no significant correlations found between fast spindle density and change in memory for 2PRES or 4PRES items (**Figure 4A**). The magnitude of the fast spindle - 1PRES correlation, averaged across the significant electrode cluster, was significantly larger than the 2PRES (*r* = -.10, *p* =.96) and 4PRES (*r* = -.11, *p* = .29) conditions (1PRES vs 2PRES, *z* = 2.40, *p* = .008; 1PRES vs 4PRES, *z* = 2.38, *p* = .009).

**Figure 4.**
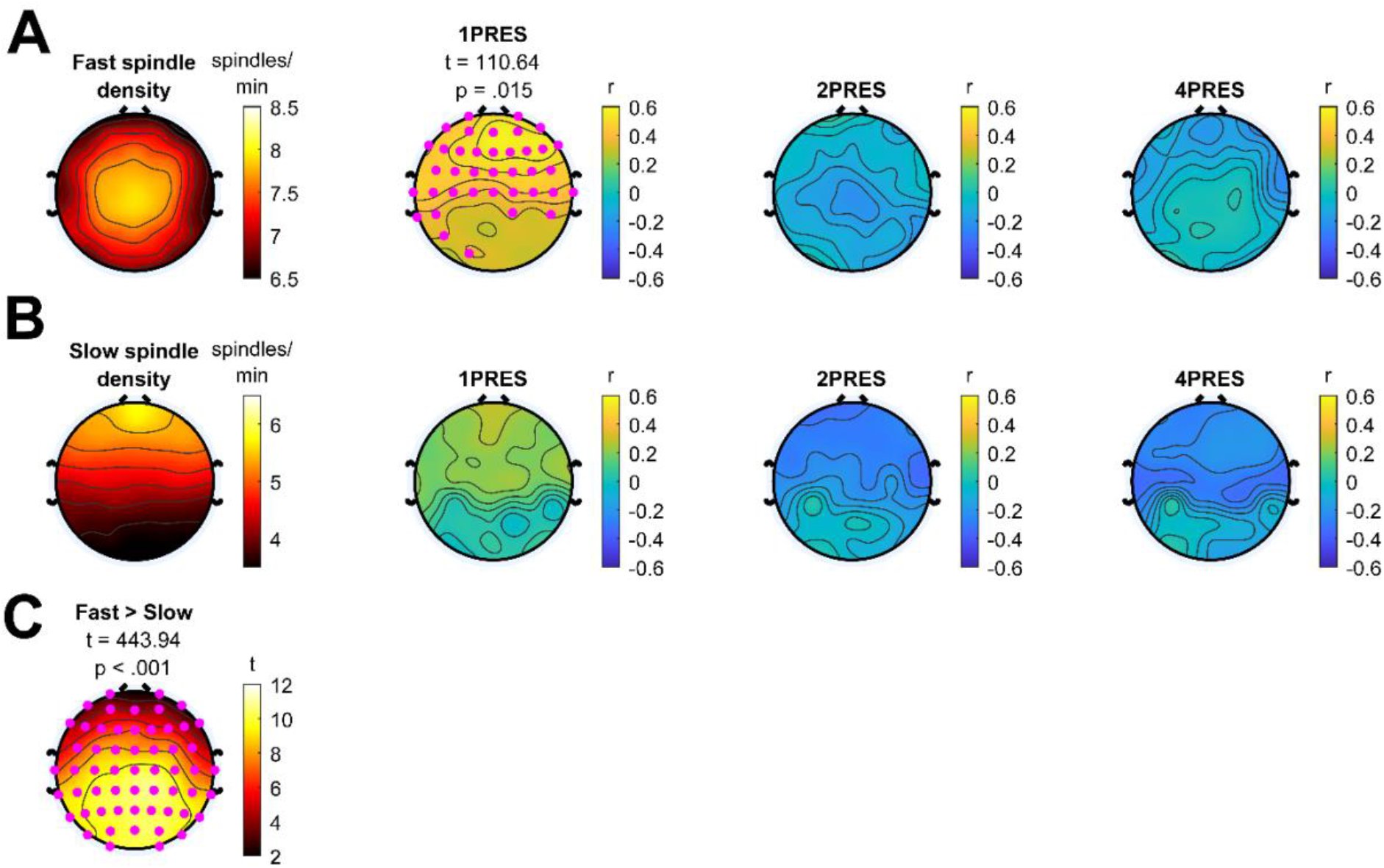
Sleep spindles during NREM sleep. **A** – Fast sleep spindles. Fast spindle density (spindles/min) during NREM sleep shown in column 1. Columns 2-4 show correlations between fast spindle density and change in recall for 1PRES, 2PRES, and 4PRES items respectively. Cluster statistics presented above topographies. Pink dots indicate significant electrodes in cluster.**B** – Slow sleep spindles. Slow spindle density (spindles/min) during NREM sleep shown in column 2. Columns 2-4 show correlations between slow spindle density and change in recall for 1PRES, 2PRES, and 4PRES items respectively. **C** – Topography of t-values indicating differences between fast and slow spindle density. Cluster statistics presented above topography. Pink dots indicate significant electrodes in cluster.

**Figure 5.**
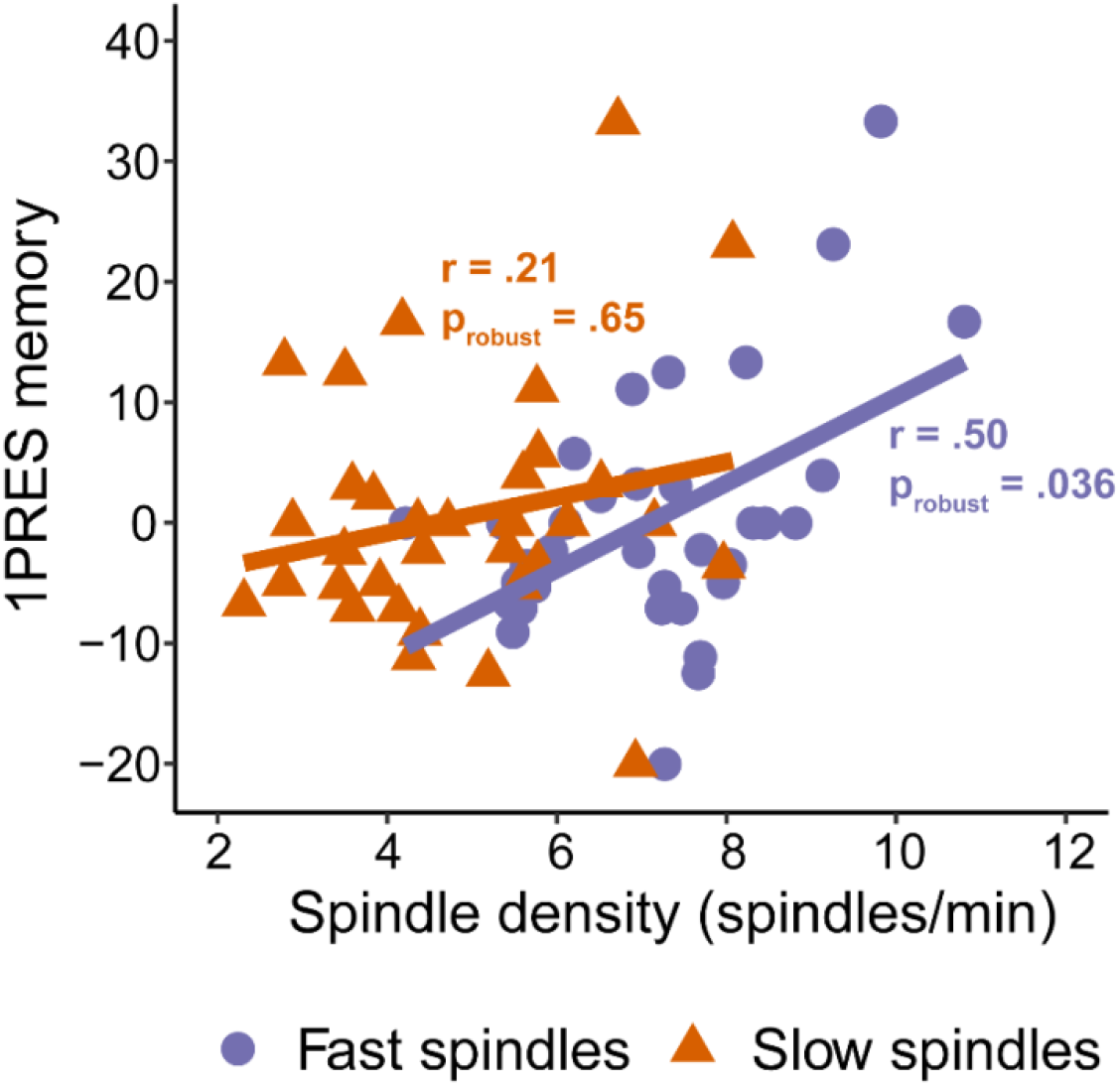
Fast and slow spindles show opposite associations with 1PRES memory. Spindle density derived by averaging significant electrodes in cluster (see topographies). P values indicate significance of a robust regression, used to minimize the influence of outliers.

For slow spindle density, we did not observe any significant correlations with change in recall for any item condition, although there was a non-significant negative association between slow spindle density and memory for 2PRES and 4PRES items (**Figure 4**). Although only fast spindle density was significantly associated with memory for the 1PRES items, when we directly compared the magnitude of the association fast and slow spindles had with 1PRES memory (again averaging together all electrodes in the significant cluster), they did not differ significantly at the .05 level (*z* = 1.41, *p* = .078). There were no significant correlations (cluster-corrected) between either fast or sleep spindles and immediate recall performance (all *p*’s > .08). This suggests that the effects of spindle density are related primarily to the consolidation that occurred during sleep. In summary, these analyses show that sleep spindles are preferentially correlated with change in recall for 1PRES items, and this association appears to be strongest for fast spindles.

### Slow oscillation-spindle coupling

A key tenet of the active systems consolidation theory is that memories are reactivated and thus consolidated through the precise coupling of hippocampal sharp-wave ripples, thalamocortical sleep spindles, and cortical slow oscillations (Klinzing et al., 2019). Prior studies have indicated that slow oscillation-spindle coupling at the scalp EEG level is correlated with memory consolidation (Demanuele et al., 2016; Mikutta et al., 2019; Muehlroth et al., 2019). It has also previously been shown that fast and slow spindles couple at different phases of the slow oscillation, which may in part reflect a functional difference between these two spindle types (Mölle et al, 2011; Klinzing et al, 2016; Cox et al., 2018).

Slow oscillation-spindle coupling for fast and slow spindles is shown in **Figure 6**. Across the whole scalp, 20% (SD = 4%) of fast spindles and 21% (SD = 4%) of slow spindles were coupled to a slow oscillation. All participants exhibited coupling of both fast and slow spindles at all electrode sites. Coupled spindle density for both fast and slow spindles is displayed in **Figure 6A**. Observed coupling rates significantly exceeded what would be expected by chance in the majority of participants (70%, SD = 8%), averaging across all electrode sites; **Figure 6A**). Average coupling phases of slow and fast spindles are displayed in **Figure 6B**. For fast spindles, Rayleigh tests of non-uniformity were significant at all electrodes sites (all *p*_adj_ < .001), suggesting the coupling of fast spindles to slow oscillations were non-uniform. In line with previous work, fast spindles were preferentially coupled to the rising slope of the slow oscillation. There was topographical variance such that, when moving from anterior to posterior, the preferred coupling phase shifted to occurring earlier in posterior channels compared to frontal channels. Slow spindles showed a different pattern of coupling to the slow oscillation, and preferentially coupled on the downward slope from the peak to the trough. For slow spindle coupling, non-uniformity tests were only significant in the frontal half of the electrodes (**Figure 6B**). When averaging across electrodes showing significant non-uniformity, the difference in coupling phase between fast (M = −41.72°, SD = 34.19) and slow spindles (M =110.92°, SD = 56.81°) was highly significant (*F* = 75.74,*p* < .001). Visualization of the temporal dynamics underlying fast and slow spindle coupling are shown in **Figure 6C**.

**Figure 6.**
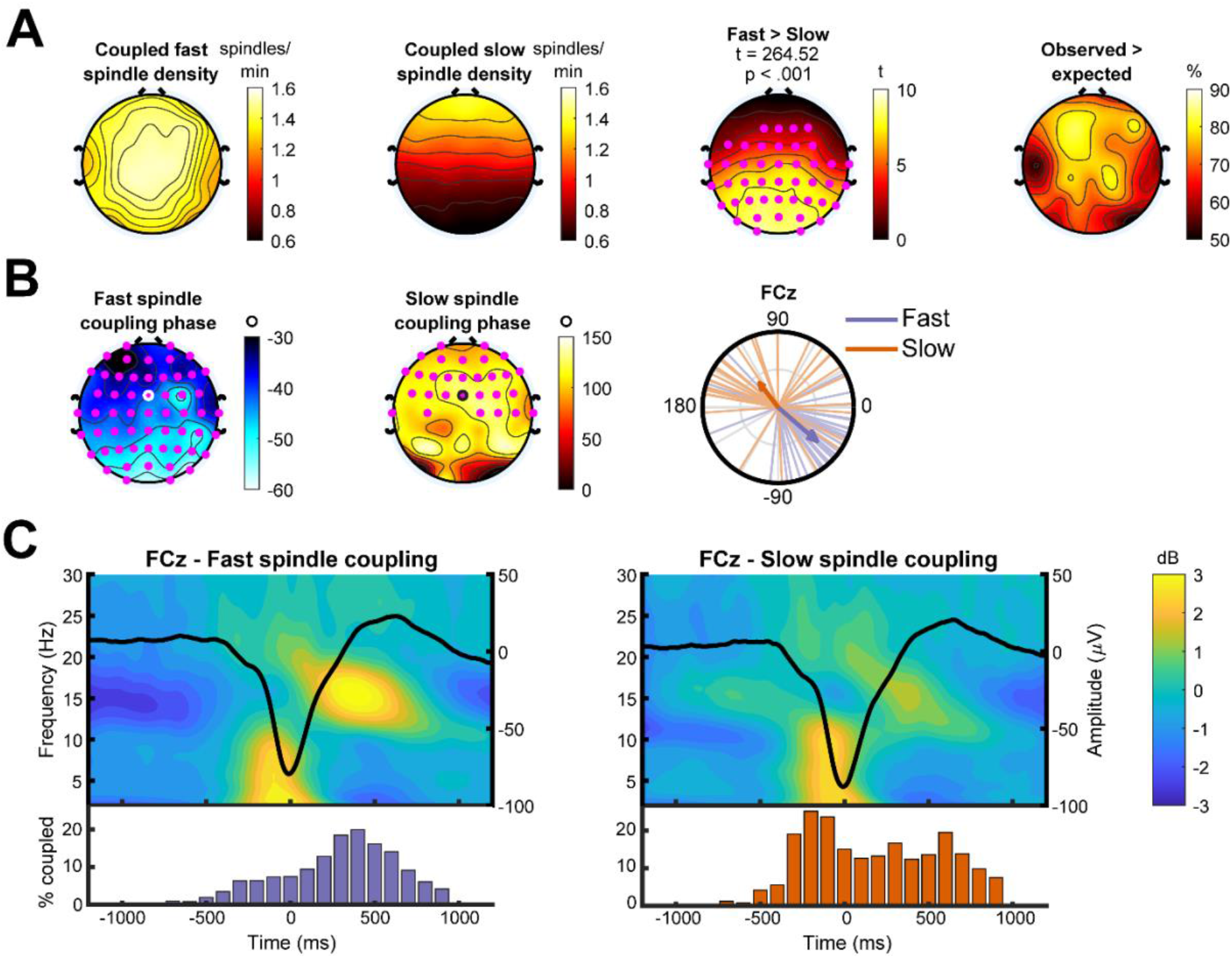
Slow oscillation-spindle coupling. **A** – Topography of coupled fast (first column) and slow (second column) spindle density, and the difference between them (third column). Cluster statistics presented above topography. Pink dots indicate significant electrodes in cluster. The percentage of participants who exhibited a significantly higher rate of coupling than would be expected by chance is shown in the far-right column. **B** – Coupling phase. Columns 1 and 2 shows preferred coupling phase of fast and slow spindles at each electrode. Pink dots indicate electrodes exhibiting significant nonuniformity (Rayleigh tests) at *p* < .05 (adjusted for multiple comparisons using the False Discovery Rate). Column 3 shows a circular phase plot displaying phase distributions of fast and slow spindles across participants at electrode FCz (ringed electrode in topographies). Each line indicates the preferred coupling phase of individual participants. The direction of the arrow indicates the average phase across participants, with the length of the arrow indicating the coupling strength across participants. A coupling of phase of 0° indicates preferential coupling of spindles at the positive peak of the slow oscillation. A coupling phase of 180° indicates preferential spindle coupling at the negative trough of the slow oscillation. **C** – Temporal dynamics of slow oscillation-spindle coupling events. In the left-hand figure, the top plot shows the average time-frequency response of all slow oscillations coupled to a fast spindle (−1200 – 1200ms, centered on the trough of the slow oscillation), with the time-domain averaged slow oscillation overlaid. Underneath, a histogram indicating the distribution of coupled fast spindles, displayed as % of coupled fast spindles, averaged across participants, binned into 100ms intervals. The right-hand plot displays the same information, but for slow spindle coupling. All analyses performed on electrode FCz (ringed electrode in topographies of panel B).

We then sought to investigate how slow oscillation-spindle coupling was associated with change in recall across the delay period. Given our findings above (**Figures 4–5**), we focused our attention primarily on fast spindles, and on the association between coupling and 1PRES change in recall. Both coupled (cluster *t* = 132.10, *p* = .003) and uncoupled (cluster *t* = 83.37, *p* = .030) fast spindle density showed positive correlations with 1PRES change in recall (**Figure 7A**; average within-cluster correlation: coupled fast spindles (*r* = .52, *p* = .002), uncoupled fast spindles (*r* = .45, *p* = .008).

**Figure 7.**
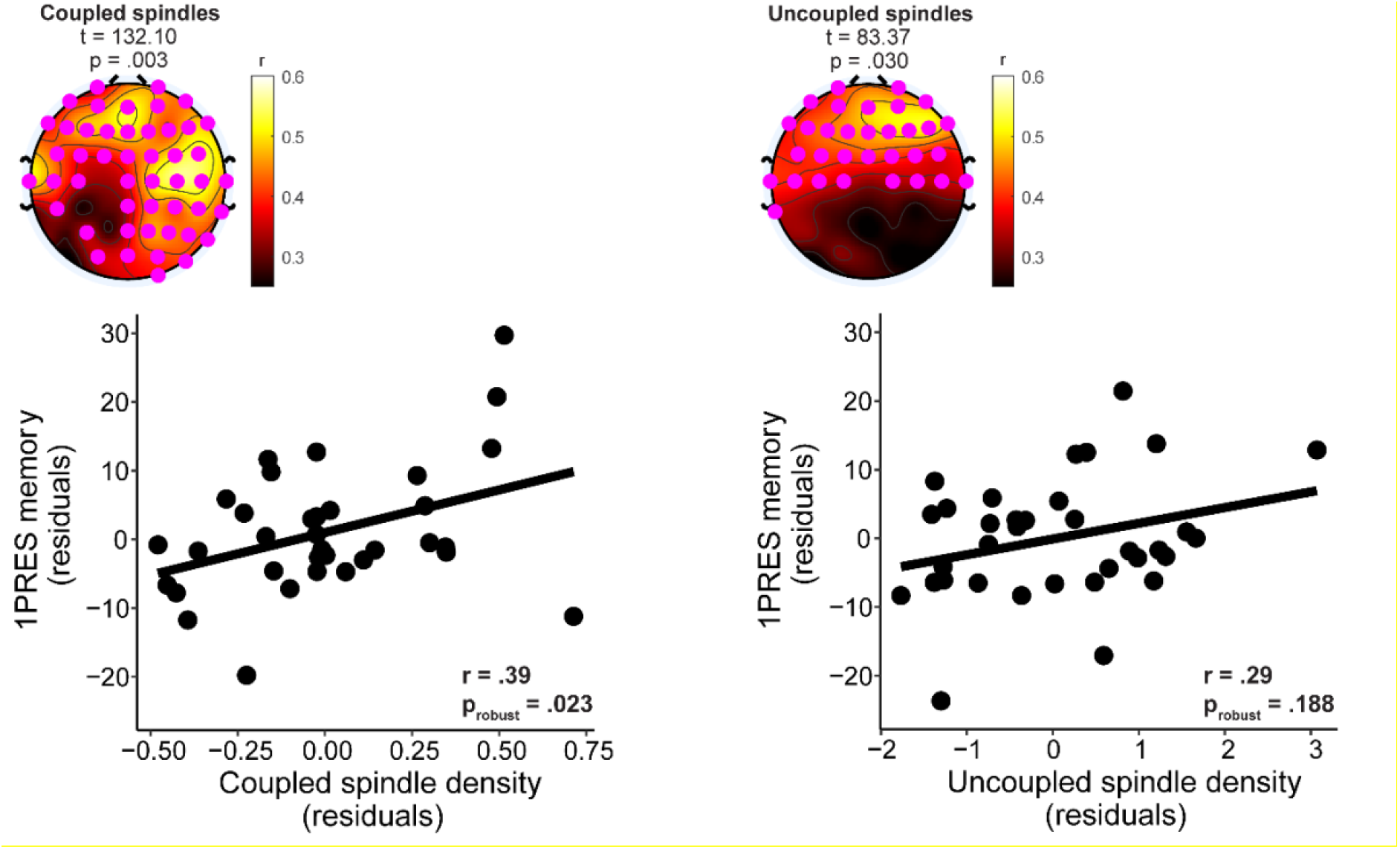
Fast spindle coupling and memory consolidation of 1PRES items. Top row, univariate correlations between fast coupled spindle density (left plot) and fast uncoupled spindle density (right plot) with 1PRES change in recall. Cluster statistics presented above topographies. Pink dots indicate significant electrodes in cluster. Bottom row, partial robust regression plots. Left plot visualizes the relationship between coupled fast spindle density and 1PRES change in recall, after accounting for uncoupled spindles. The right plot visualizes the relationship between uncoupled fast spindle density and 1PRES change in recall, after accounting for coupled spindles.

To determine whether coupled spindles are associated with memory *independently* of uncoupled spindles, we ran a robust multiple linear regression predicting 1PRES change in recall from coupled and uncoupled spindle densities. Coupled spindle density estimates were derived by averaging spindle density across all electrodes in the significant clusters for coupled and uncoupled spindles respectively (**Figure 7A**). The overall model was significant (*F* (1, 31) = 8.88, *p* < .001, adjusted R^2^ = .25), but only coupled spindle density significantly predicted 1PRES memory; β [95% CI] = 4.22 [0.72, 7.72], *p* = .020 (**Figure 7B**). Uncoupled spindle density did not predict 1PRES memory independently of coupled spindle density (β [95% CI] = 2.16 [−1.33, 5.66], *p* = .22; **Figure 7B**). These results suggest that the density of coupled spindles were uniquely related to the consolidation of 1PRES items independently of uncoupled spindles. There were no significant circular-linear correlations between fast spindle coupling phase and the change in recall for 1PRES items at an FDR adjusted *p* < .05 level (all *r* < .44).

Supplementary post-hoc tests found no significant clusters for coupled or uncoupled fast spindles when correlated with change in recall for either 2PRES or 4PRES items, nor were any significant clusters found for any correlations between coupled or uncoupled slow spindles and change in recall for any encoding strength condition.

### Subjective sleep measures

Subjective sleep variables are shown in **Table 3.**At the start of the second session, the sleep group reported feeling significantly more refreshed than the wake group (*t* (52) = 2.12, *p* = .039, *d* = 0.61). However, the change in recall between the second and first session (the key dependent variable of this study), was not associated with the subjective feeling of being refreshed at the second session in either group (sleep: *r* = .13, *p* = .45; wake: *r* = .05, *p* = .84). There were no other differences between the groups on any other of the subjective measures.

**Table 3.** Subjective sleep variables

|  | Sleep<br>M (SD) | Wake<br>M (SD) | sig | d |
| --- | --- | --- | --- | --- |
| PSQI global score | 4.6 (2.6) | 4.3 (2.7) | .64 | 0.14 |
| 3-night log sleep onset (min) | 16.8 (9.7) | 18.7 (18.2) | .63 | 0.14 |
| 3-night log hours asleep | 7.4 (0.8) | 7.7 (0.7) | .13 | 0.44 |
| 3-night log sleep quality | 1.86 (0.54) | 1.96 (0.48) | .50 | 0.20 |
| Session 1: Concentration | 73.4 (17.2) | 67.7 (21.4) | .30 | 0.30 |
| Session 1: Refreshed | 63.1 (24.5) | 52.4 (23.8) | .13 | 0.44 |
| Session 1: Sleepiness | 2.53 (0.74) | 2.50 (0.99) | .91 | 0.03 |
| Session 2: Concentration | 79.9 (14.4) | 73.4 (19.7) | .18 | 0.40 |
| Session 2: Refreshed | 75.4 (20.4) | 61.7 (25.8) | .039 | 0.61 |
| Session 2: Sleepiness | 2.0 (0.83) | 2.3 (1.03) | .21 | 0.37 |

*Note.* Sig = *p* value for between-groups t-test. PSQI = Pittsburgh sleep quality index (theoretical range = 0-21, a higher score = worse sleep quality). 3-night sleep log quality theoretical range = 1 - 4, a higher score = worse sleep quality. Concentration and refreshed items theoretical range = 0 - 100, a higher score = better concentration / more refreshed. Sleepiness item theoretical range = 1-8, a higher score = greater subjective sleepiness.

### Discussion

Here, we set out to investigate how encoding strength influences the consolidation of memories over a six-hour delay with or without a daytime nap, and what the sleep EEG oscillatory correlates of sleep-based consolidation were. With regards to the first aim, we found that over a wake-only delay, there was significantly more forgetting of weakly encoded memories compared to strong ones. It is likely that after just a single presentation during encoding, the neural representations of these memories are weaker and less stable compared to memories for items that were presented multiple times (Xue et al., 2010; Lu et al., 2015). Here, it appeared that those weak memory traces fade from memory faster than stronger, more durable traces.

When participants were able to nap shortly after learning, however, we did not see this pattern. Instead, the rate of forgetting was the same across all three encoding strengths. When forgetting rates were compared between the nap and the wake group, the only significant difference was more forgetting of weak items in the group that did not nap. As such, it appears that sleep and wake differentially affect the consolidation of memories of different strengths. Over a period of sleep, weakly encoded memories received a consolidation benefit, whilst those same memories deteriorated the fastest over a wake delay. It is important to note that these nap-related benefits were seen after four hours elapsed between the end of the nap and the delayed test, suggesting that the nap facilitated a stabilizing effect on the weak memories such that they were resilient to forgetting across the post-nap delay. Future studies should investigate whether these benefits of a daytime nap would be seen over longer delay intervals.

The results of this nap study are broadly comparable to our previous work, where we used the same method to induce different encoding strengths, but focused on nocturnal sleep and a 12 or 24hr delay interval (Denis et al., 2020). In that study, we found that across a 12hr wake delay, forgetting of weak memories was significantly greater than strong memories. Interestingly, the ~30% forgetting of weak memories across a 12-hour delay seen in that study was approximately half that seen in the current study over a 6-hour delay (~13%), suggesting a linear increase in forgetting of weak memories with time spent awake. On the other hand, across a 12-hour delay filled with nocturnal sleep, forgetting of weak memories was attenuated and appeared to be stabilized over a subsequent 12 hours of wakefulness, to a degree similar to that seen after the 6hr delay in the present study (Denis et al., 2020).

In our previous study, we found that weakly encoded items (manipulated by the number of exposures) were only consolidated when the items had been successfully visualized during encoding. We were unable to investigate the role of visualization directly in the present study, as participants reported being able to visualize a far larger percentage (78%) of items than in that study (54%). The present study used a 2,000ms presentation time, compared to 1,500ms previously, which may have made it more likely that a word pair would be visualized (Denis et al., 2020).

Our second question pertained to the electrophysiological or sleep correlates of memory consolidation across the nap. We did not find the amount of time spent in any particular sleep stage was associated with memory consolidation, despite earlier studies finding a link between word-pair memory and SWS (Plihal and Born, 1997). We did, however, find that sleep spindles were associated with memory consolidation for the weakly encoded information. However, this was dependent on spindle type. For fast spindles, higher density (spindles/min) was associated with less forgetting of weakly encoded memories, whilst slow spindles were not associated with consolidation of any item category (and in fact showed a non-significant negative association with more strongly encoded items). This dissociation adds to the growing body of work suggesting differential functional roles for fast and slow spindles with relation to memory consolidation (Fernandez and Luthi, 2020).

The active systems consolidation hypothesis emphasizes the importance of coupling between key cortical and subcortical oscillations, and indeed a number of prior studies have shown the degree of coupling to be associated with sleep-based consolidation (Mölle et al., 2009; Niknazar et al., 2015; Demanuele et al., 2016; Muehlroth et al., 2019). When considering fast spindles, we found that coupled spindle density predicted the consolidation of weakly encoded memories, even after controlling for the effect of uncoupled spindle density. As such, our data adds further support to the idea that slow oscillation-spindle coupling events are critically involved in memory consolidation processes occurring during sleep. Although we only found coupled spindles to independently predict memory, we do note that univariate correlations did show a significant positive correlation between uncoupled spindles and memory. This indicates that uncoupled spindles do play some role in memory consolidation processes, albeit potentially not as strong a role as coupled spindles. This finding warrants future work to better delineate the origins and functions of sleep spindles at either are or are not coupled to the slow oscillation.

Sleep spindles have been shown to relate to cognitive ability more generally (Schabus et al., 2008; Ujma, 2018), and some have shown that increased spindle activity is related to higher memory recall both before and after sleep (Gais et al., 2002). We did not find sleep spindles to be associated with memory at the immediate test, but rather only to the change in recall across the delay period. That suggests, in this dataset at least, that spindle density is more related to consolidation processes. In a recent meta-analysis, it was concluded that spindle amplitude and duration, rather than density, were positively correlated with general cognitive ability (Ujma, 2018). As such, different spindle properties may relate differently to memory consolidation mechanisms and more general cognitive aptitude.

We found that fast spindle density was only associated with consolidation of weakly encoded items and was not related to the consolidation of the more strongly encoded information. This mirrors our behavioral finding, where the only significant difference between the nap and the wake group was for the weakly encoded items. It is believed that slow oscillation-spindle (and hippocampal ripple) coupling facilitates the reactivation of memory traces in the hippocampus, which promotes the strengthening of memory traces in hippocampal-neocortical circuits (Takashima et al., 2006; Payne and Kensinger, 2011; Latchoumane et al., 2017; Zhang et al., 2018; Schreiner et al., 2020). In this study, it is possible that only the weakly encoded items needed to undergo this strengthening during sleep, as the more strongly encoded items were already well consolidated before the retention interval occurred. Other research has shown that extremely well-formed memories do not benefit from subsequent sleep (Himmer et al., 2017), and repeated learning cycles can induce a ‘fast’ consolidation process (Brodt et al., 2018).

It is important to note that our findings are correlational in nature. Other work has shown that it is possible to enhance spindle activity pharmacologically to alter memory function, (Kaestner et al., 2013; Niknazar et al., 2015), though potentially at the cost of disrupting slow oscillation-spindle coupling processes (Mylonas et al., 2020). Furthermore, commonly used pharmacological agents such as zolpidem and eszopiclone have been shown to induce widespread alterations in the sleep EEG, and as such do not solely target changes in sleep spindles (Aeschbach et al., 1994; Monti et al., 2000; Struyk et al., 2016). Acoustic stimulation has also been used to increase the number of sleep spindles (Antony and Paller, 2017). Future work should further embrace the use of these methods in determining the memory function of sleep spindles.

Sleep benefits the consolidation of memories, a process which is facilitated by the coupling of slow oscillations and sleep spindles during NREM sleep (Demanuele et al., 2016; Helfrich et al., 2018; Klinzing et al., 2019; Mikutta et al., 2019). This benefit is not uniform, with some memories being preferentially consolidated over others (Payne, 2011; Stickgold and Walker, 2013). Here, we show that sleep spindles preferentially consolidate memories that are weakly encoded initially. This may be due to these memories being in the most need of being consolidated, compared to stronger memories that are already adequately strengthened before sleep. As such, these results demonstrate initial encoding strength to be an important boundary condition in determining when sleep-based memory consolidation is likely to occur.

## Acknowledgements

This work was supported by a National Institutes of Health grant (R01-MH48832), awarded to RS. JDP is supported by a National Science Foundation grant (BCS-2001025). We would like to thank the anonymous reviewers for their thoughtful and constructive suggestions.

## References

Abel M, Haller V, Köck H, Pötschke S, Heib D, Schabus M, Bäuml K-HT (2019) Sleep reduces the testing effect-But not after corrective feedback and prolonged retention interval. J Exp Psychol Learn Mem Cogn 45:272–287.

Aeschbach D, Dijk D-J, Trachsel L, Brunner DP, Borbély AA (1994) Dynamics of Slow-Wave Activity and Spindle Frequency Activity in the Human Sleep EEG: Effect of Midazolam and Zopiclone. Neuropsychopharmacology 11:237–244.

Andrade J, May J, Deeprose C, Baugh S-J, Ganis G (2014) Assessing vividness of mental imagery: The Plymouth Sensory Imagery Questionnaire. Br J Psychol 105:547–563.

Antony JW, Paller KA (2017) Using Oscillating Sounds to Manipulate Sleep Spindles. Sleep 40:zsw068.

Baena D, Cantero JL, Fuentemilla L, Atienza M (2020) Weakly encoded memories due to acute sleep restriction can be rescued after one night of recovery sleep. Scientific Reports 10:1449.

Bates D, Mächler M, Bolker B, Walker S (2015) Fitting linear mixed-effects models using lme4. J Stat Softw 67:1–48.

Bäuml K-HT, Holterman C, Abel M (2014) Sleep can reduce the testing effect: It enhances recall of restudied items but can leave recall of retrieved items unaffected. Journal of Experimental Psychology: Learning, Memory, and Cognition 40:1568–1581.

Berens P (2009) CircStat: A Matlab toolbox for circular statistics. Journal of statistical software 31.

Brodt S, Gais S, Beck J, Erb M, Scheffler K, Schönauer M (2018) Fast track to the neocortex: A memory engram in the posterior parietal cortex. Science 362:1045–1048.

Buysse DJ, Reynolds C, Monk T, Berman S, Kupfer D (1989) The Pittsburgh Sleep Quality Index (PSQI): A new instrument for psychiatric practice and research. Psychiatry Research 28:193–213.

Cohen MX (2017) Comparison of linear spatial filters for identifying oscillatory activity in multichannel data. Journal of Neuroscience Methods 278:1–12.

Cowan E, Liu A, Henin S, Kothare S, Devinsky O, Davachi L (2020) Sleep Spindles Promote the Restructuring of Memory Representations in Ventromedial Prefrontal Cortex through Enhanced Hippocampal–Cortical Functional Connectivity. J Neurosci 40:1909–1919.

Cox R, Mylonas DS, Manoach DS, Stickgold R (2018) Large-scale structure and individual fingerprints of locally coupled sleep oscillations. Sleep 41:zsy175.

Cox R, Schapiro AC, Manoach DS, Stickgold R (2017) Individual differences in frequency and topography of slow and fast sleep spindles. Front Hum Neurosci 11:433.

Cox R, van Driel J, de Boer M, Talamini LM (2014) Slow oscillations during sleep coordinate interregional communication in cortical networks. J Neurosci 34:16890–16901.

Creery JD, Oudiette D, Antony JW, Paller KA (2015) Targeted Memory Reactivation during Sleep Depends on Prior Learning. Sleep 38:755–763.

Cross ZR, Helfrich RF, Kohler MJ, Corcoran AW, Coussens S, Zou-Williams L, Schlesewsky M, Gaskell MG, Knight RT, Bornkessel-Schlesewsky I (2020) Slow wave-spindle coupling during sleep predicts language learning and associated oscillatory activity. bioRxiv:2020.02.13.948539.

Delorme A, Makeig S (2004) EEGLAB: an open source toolbox for analysis of single-trial EEG dynamics including independent component analysis. J Neurosci Methods 134:9–21.

Demanuele C, Bartsch U, Baran B, Khan S, Vangel MG, Cox R, Hämäläinen M, Jones MW, Stickgold R, Manoach DS (2016) Coordination of Slow Waves with Sleep Spindles Predicts Sleep-Dependent Memory Consolidation in Schizophrenia. Sleep 40:zsw013.

Demanuele C, James CJ, Sonuga-Barke EJ (2007) Distinguishing low frequency oscillations within the 1/f spectral behaviour of electromagnetic brain signals. Behavioral and Brain Functions 3:62.

Denis D, Schapiro AC, Poskanzer C, Bursal V, Charon L, Morgan A, Stickgold R (2020) The roles of item exposure and visualization success in the consolidation of memories across wake and sleep. Learn Mem 27:451–456.

Diekelmann S, Wilhelm I, Born J (2009) The whats and whens of sleep-dependent memory consolidation. Sleep Med Rev 13:309–321.

Drosopoulos S, Schulze C, Fischer S, Born J (2007) Sleep’s function in the spontaneous recovery and consolidation of memories. J Exp Psychol Gen 136:169–183.

Fernandez LMJ, Luthi A (2020) Sleep Spindles: Mechanisms and Functions. Physiol Rev 100:805–868.

Fischer S, Born J (2009) Anticipated reward enhances offline learning during sleep. J Exp Psychol Learn Mem Cogn 35:1586–1593.

Fisher RA (1925) Statistical Methods for Research Workers. Edinburgh, Scotland: Oliver and Boyd. Available at: http://psychclassics.yorku.ca. [Accessed June 2, 2020].

Gais S, Mölle M, Helms K, Born J (2002) Learning-Dependent Increases in Sleep Spindle Density. J Neurosci 22:6830–6834.

Gilson M, Tauste Campo A, Chen X, Thiele A, Deco G (2017) Nonparametric test for connectivity detection in multivariate autoregressive networks and application to multiunit activity data. Network Neuroscience 1:357–380.

Helfrich RF, Mander BA, Jagust WJ, Knight RT, Walker MP (2018) Old Brains Come Uncoupled in Sleep: Slow Wave-Spindle Synchrony, Brain Atrophy, and Forgetting. Neuron 97:221–230.

Himmer L, Müller E, Gais S, Schönauer M (2017) Sleep-mediated memory consolidation depends on the level of integration at encoding. Neurobiol Learn Mem 137:101–106.

Hoddes E, Dement W, Zarcone V (1972) The development and use of the stanford sleepiness scale (SSS). Psychophysiol 9:150.

Iber C, Ancoli-Israel S, Chesson A, Quan SF (2007) The AASM Manual for the Scoring of Sleep and Associated Events: Rules, Terminology and Technical Specification. Westchester, IL: Americian Academy of Sleep Medicine.

Kaestner EJ, Wixted JT, Mednick SC (2013) Pharmacologically Increasing Sleep Spindles Enhances Recognition for Negative and High-arousal Memories. Journal of Cognitive Neuroscience 25:1597–1610.

Kim SY, Payne JD (2020) Neural correlates of sleep, stress, and selective memory consolidation. Current Opinion in Behavioral Sciences 33:57–64.

Kleiner M, Brainard D, Pelli D (2007) What’s new in Psychtoolbox-3. Perception 36:ECVP-Abstract Supplement.

Klinzing JG, Mölle M, Weber F, Supp G, Hipp JF, Engel AK, Born J (2016) Spindle activity phase-locked to sleep slow oscillations. NeuroImage 134:607–616.

Klinzing JG, Niethard N, Born J (2019) Mechanisms of systems memory consolidation during sleep. Nat Neurosci 22:1598–1610.

Kurz E-M, Conzelmann A, Barth GM, Renner TJ, Zinke K, Born J (2020) How Do Children with Autism Spectrum Disorder Form Gist Memory During Sleep? – A Study of Slow Oscillation-Spindle Coupling. Sleep Available at: https://doi.org/10.1093/sleep/zsaa290 [Accessed January 3, 2021].

Latchoumane C-FV, Ngo H-VV, Born J, Shin H-S (2017) Thalamic Spindles Promote Memory Formation during Sleep through Triple Phase-Locking of Cortical, Thalamic, and Hippocampal Rhythms. Neuron 95:424–435.

Lenth R (2018) emmeans: Estimated Marginal Means, aka Least-Squares Means. R. Available at: https://CRAN.R-project.org/package=emmeans [Accessed April 10, 2019].

Lo JC, Dijk D-J, Groeger JA (2014) Comparing the Effects of Nocturnal Sleep and Daytime Napping on Declarative Memory Consolidation. PLOS ONE 9:e108100.

Lu Y, Wang C, Chen C, Xue G (2015) Spatiotemporal Neural Pattern Similarity Supports Episodic Memory. Current Biology 25:780–785.

Lustenberger C, Wehrle F, Tüshaus L, Achermann P, Huber R (2015) The Multidimensional Aspects of Sleep Spindles and Their Relationship to Word-Pair Memory Consolidation. Sleep 38:1093–1103.

Marks DF (1973) Visual Imagery Differences in the Recall of Pictures. British Journal of Psychology 64:17–24.

Mednick S, Nakayama K, Stickgold R (2003) Sleep-dependent learning: a nap is as good as a night. Nature Neuroscience 6:697–698.

Meng X, Rosenthal R, Rubin DB (1992) Comparing correlated correlation coefficients. Psychological Bulletin 111:172–175.

Mikutta C, Feige B, Maier JG, Hertenstein E, Holz J, Riemann D, Nissen C (2019) Phase-amplitude coupling of sleep slow oscillatory and spindle activity correlates with overnight memory consolidation. Journal of Sleep Research 28:e12835.

Mölle M, Bergmann TO, Marshall L, Born J (2011) Fast and Slow Spindles during the Sleep Slow Oscillation: Disparate Coalescence and Engagement in Memory Processing. Sleep 34:1411–1421.

Mölle M, Eschenko O, Gais S, Sara SJ, Born J (2009) The influence of learning on sleep slow oscillations and associated spindles and ripples in humans and rats. Eur J Neurosci 29:1071–1081.

Monti JM, Alvariño F, Monti D (2000) Conventional and power spectrum analysis of the effects of zolpidem on sleep EEG in patients with chronic primary insomnia. Sleep 23:1075–1084.

Muehlroth BE, Sander MC, Fandakova Y, Grandy TH, Rasch B, Lee Shing Y, Werkle-Bergner M (2020) Memory quality modulates the effect of aging on memory consolidation during sleep: Reduced maintenance but intact gain. NeuroImage 209:116490.

Muehlroth BE, Sander MC, Fandakova Y, Grandy TH, Rasch B, Shing YL, Werkle-Bergner M (2019) Precise Slow Oscillation-Spindle Coupling Promotes Memory Consolidation in Younger and Older Adults. Sci Rep 9:1940.

Mylonas D, Baran B, Demanuele C, Cox R, Vuper TC, Seicol BJ, Fowler RA, Correll D, Parr E, Callahan CE, Morgan A, Henderson D, Vangel M, Stickgold R, Manoach DS (2020) The effects of eszopiclone on sleep spindles and memory consolidation in schizophrenia: a randomized clinical trial. Neuropsychopharmacology:1–12.

Mylonas D, Tocci C, Coon WG, Baran B, Kohnke EJ, Zhu L, Vangel MG, Stickgold R, Manoach DS (2019) Naps reliably estimate nocturnal sleep spindle density in health and schizophrenia. Journal of Sleep Research 29:e12968.

Niknazar M, Krishnan GP, Bazhenov M, Mednick SC (2015) Coupling of Thalamocortical Sleep Oscillations Are Important for Memory Consolidation in Humans. PLOS ONE 10:e0144720.

Oostenveld R, Fries P, Maris E, Schoffelen J-M (2011) FieldTrip: Open source software for advanced analysis of MEG, EEG, and invasive electrophysiological data. Comput Intell Neurosci 2011:156869.

Otsu N (1979) A Threshold Selection Method from Gray-Level Histograms. IEEE Transactions on Systems, Man, and Cybernetics 9:62–66.

Payne JD (2011) Sleep on it!: stabilizing and transforming memories during sleep. Nature Neuroscience 14:272–274.

Payne JD, Chambers AM, Kensinger EA (2012) Sleep promotes lasting changes in selective memory for emotional scenes. Front Integr Neurosci 6:108.

Payne JD, Ellenbogen JM, Walker MP, Stickgold R (2008a) The role of sleep in memory consolidation. In: Learning and memory: A comprehensive reference: Vol. 2. Cognitive psychology of memory, pp 663–685. Oxford, England: Elsevier.

Payne JD, Kensinger EA (2011) Sleep leads to changes in the emotional memory trace: evidence from FMRI. J Cogn Neurosci 23:1285–1297.

Payne JD, Kensinger EA (2018) Stress, sleep, and the selective consolidation of emotional memories. Current Opinion in Behavioral Sciences 19:36–43.

Payne JD, Kensinger EA, Wamsley EJ, Spreng RN, Alger SE, Gibler K, Schacter DL, Stickgold R (2015) Napping and the selective consolidation of negative aspects of scenes. Emotion 15:176–186.

Payne JD, Schacter DL, Propper RE, Huang L-W, Wamsley EJ, Tucker MA, Walker MP, Stickgold R (2009) The role of sleep in false memory formation. Neurobiology of Learning and Memory 92:327–334.

Payne JD, Stickgold R, Swanberg K, Kensinger EA (2008b) Sleep preferentially enhances memory for emotional components of scenes. Psychol Sci 19:781–788.

Plihal W, Born J (1997) Effects of early and late nocturnal sleep on declarative and procedural memory. J Cogn Neurosci 9:534–547.

R Core Team (2018) R: A language and environment for statistical computing. Vienna, Austrua: R Foundation for Statistical Computing. Available at: https://www.R-project.org [Accessed February 25, 2019].

Rasch B, Born J (2013) About sleep’s role in memory. Physiol Rev 93:681–766.

Rauchs G, Feyers D, Landeau B, Bastin C, Luxen A, Maquet P, Collette F (2011) Sleep contributes to the strengthening of some memories over others, depending on hippocampal activity at learning. J Neurosci 31:2563–2568.

Roediger HL, Butler AC (2011) The critical role of retrieval practice in long-term retention. Trends Cogn Sci (Regul Ed) 15:20–27.

Schabus M, Hödlmoser K, Gruber G, Sauter C, Anderer P, Klösch G, Parapatics S, Saletu B, Klimesch W, Zeitlhofer J (2006) Sleep spindle-related activity in the human EEG and its relation to general cognitive and learning abilities. Eur J Neurosci 23:1738–1746.

Schabus M, Hoedlmoser K, Pecherstorfer T, Anderer P, Gruber G, Parapatics S, Sauter C, Kloesch G, Klimesch W, Saletu B, Zeitlhofer J (2008) Interindividual sleep spindle differences and their relation to learning-related enhancements. Brain Res 1191:127–135.

Schapiro AC, McDevitt EA, Chen L, Norman KA, Mednick SC, Rogers TT (2017) Sleep Benefits Memory for Semantic Category Structure While Preserving Exemplar-Specific Information. Sci Rep 7:14869.

Schoch SF, Cordi MJ, Rasch B (2017) Modulating influences of memory strength and sensitivity of the retrieval test on the detectability of the sleep consolidation effect. Neurobiol Learn Mem 145:181–189.

Schreiner T, Petzka M, Staudigl T, Staresina BP (2020) Endogenous memory reactivation during sleep in humans is clocked by slow oscillation-spindle complexes. bioRxiv:2020.09.16.299545.

Sleigh JW, Steyn-Ross DA, Steyn-Ross ML, Williams ML, Smith P (2001) Comparison of changes in electroencephalographic measures during induction of general anaesthesia: influence of the gamma frequency band and electromyogram signal. British Journal of Anaesthesia 86:50–58.

Spaak E (2020) Meng’s Z-test for correlated correlation coefficients. MATLAB Central File Exchange Available at: https://www.mathworks.com/matlabcentral/fileexchange/37867-meng-s-z-test-for-correlated-correlation-coefficients [Accessed January 8, 2020].

Staresina BP, Bergmann TO, Bonnefond M, van der Meij R, Jensen O, Deuker L, Elger CE, Axmacher N, Fell J (2015) Hierarchical nesting of slow oscillations, spindles and ripples in the human hippocampus during sleep. Nature Neuroscience 18:1679–1686.

Stickgold R (2005) Sleep-dependent memory consolidation. Nature 437:1272–1278.

Stickgold R (2009) How do I remember? Let me count the ways. Sleep Med Rev 13:305–308.

Stickgold R, Walker MP (2013) Sleep-dependent memory triage: evolving generalization through selective processing. Nat Neurosci 16:139–145.

Struyk A, Gargano C, Drexel M, Stoch SA, Svetnik V, Ma J, Mayleben D (2016) Pharmacodynamic effects of suvorexant and zolpidem on EEG during sleep in healthy subjects. European Neuropsychopharmacology 26:1649–1656.

Takashima A, Petersson KM, Rutters F, Tendolkar I, Jensen O, Zwarts MJ, McNaughton BL, Fernández G (2006) Declarative memory consolidation in humans: a prospective functional magnetic resonance imaging study. Proc Natl Acad Sci USA 103:756–761.

Tucker MA, Fishbein W (2008) Enhancement of declarative memory performance following a daytime nap is contingent on strength of initial task acquisition. Sleep 31:197–203.

Ujma PP (2018) Sleep spindles and general cognitive ability – A meta-analysis. Sleep Spindles & Cortical Up States:1–17.

van den Brink RL (2020) circt_htest.m. MATLAB Central File Exchange Available at: https://www.mathworks.com/matlabcentral/fileexchange/46349-circ_htest-m [Accessed September 28, 2020].

van den Brink RL, Wynn SC, Nieuwenhuis S (2014) Post-Error Slowing as a Consequence of Disturbed Low-Frequency Oscillatory Phase Entrainment. J Neurosci 34:11096–11105.

van Schalkwijk FJ, Sauter C, Hoedlmoser K, Heib DPJ, Klösch G, Moser D, Gruber G, Anderer P, Zeitlhofer J, Schabus M (2017) The effect of daytime napping and full-night sleep on the consolidation of declarative and procedural information. J Sleep Res 28 Available at: http://dx.doi.org/10.1111/jsr.12649 [Accessed November 19, 2018].

Velleman PF, Welsch RE (1981) Efficient Computing of Regression Diagnostics. The American Statistician 35:234–242.

Wamsley EJ, Tucker MA, Shinn AK, Ono KE, McKinley SK, Ely AV, Goff DC, Stickgold R, Manoach DS (2012) Reduced sleep spindles and spindle coherence in schizophrenia: mechanisms of impaired memory consolidation? Biol Psychiatry 71:154–161.

Wilhelm I, Diekelmann S, Molzow I, Ayoub A, Mölle M, Born J (2011) Sleep selectively enhances memory expected to be of future relevance. J Neurosci 31:1563–1569.

Wislowska M, Heib DPJ, Griessenberger H, Hoedlmoser K, Schabus M (2017) Individual baseline memory performance and its significance for sleep-dependent memory consolidation. Sleep Spindles & Cortical Up States 1:2–13.

Xue G, Dong Q, Chen C, Lu Z, Mumford JA, Poldrack RA (2010) Greater Neural Pattern Similarity Across Repetitions Is Associated with Better Memory. Science 330:97–101.

Zhang H, Fell J, Axmacher N (2018) Electrophysiological mechanisms of human memory consolidation. Nat Commun 9:4103.

